# Acoustogenetic Control of CAR T Cells via Focused Ultrasound

**DOI:** 10.1101/2020.02.18.955005

**Authors:** Yiqian Wu, Yahan Liu, Ziliang Huang, Xin Wang, Zhen Jin, Jiayi Li, Praopim Limsakul, Linshan Zhu, Molly Allen, Yijia Pan, Robert Bussell, Aaron Jacobson, Thomas Liu, Shu Chien, Yingxiao Wang

## Abstract

Optogenetics can control specific molecular events in living systems, but the penetration depth of light is typically limited at hundreds of micrometers. Focused ultrasound (FUS), on the other hand, can deliver energy safely and noninvasively into tissues at depths of centimeters. Here we have developed an acoustogenetic approach using short-pulsed FUS to remotely and directly control the genetics and cellular functions of engineered mammalian cells for therapeutic purposes. We applied this acoustogenetic approach to control chimeric antigen receptor (CAR) T cells with high spatiotemporal precision, aiming to mitigate the potentially lethal “on-target off-tumor” effects of CAR T cell therapy. We first verified the controllability of our acoustogenetic CAR T cells in recognizing and killing tumor cells *in vitro*, and then applied this approach *in vivo* to suppress tumor growth of both lymphoma and prostate cancers. The results indicate that FUS-based acoustogenetics can allow the noninvasive and remote activation, without any exogenous cofactor, of different types of CAR T cells for cancer therapeutics.

---

Optogenetics enables the control of specific molecular events and cellular functions in living systems with high spatiotemporal resolutions. However, optogenetics cannot reach deep tissues, with the penetration depth of light typically limited at micrometer to millimeter scales (*1*). Ultrasound can be focused to deliver mechanical energy safely and noninvasively into small volumes of tissue deep inside the body up to tens of centimeters (*1*). The rapidly oscillating pressure of focused ultrasound (FUS) waves and the resultant cycles of mechanical loading/unloading can lead to local heat generation in biological tissues. Aided by Magnetic Resonance Imaging (MRI) thermometry, FUS has been widely applied to clinically ablate tumors, and control drug delivery, vasodilation, neuromodulation (*2*), and transgene expression (*3–5*). Transcription factors and genetic circuits have also been engineered to convert the FUS-generated heat into genetic regulation to control microbial systems *in vivo* (*6*). However, there is a lack of general methods using FUS to control mammalian cell functions *in vivo* for therapeutic applications.

Chimeric antigen receptor (CAR) T cell therapy, where T cells are genetically programmed with redirected specificity against malignant cells, is becoming a paradigm-shifting approach for cancer treatment, especially for blood cancers (*7*). However, major challenges remain for solid tumors before CAR-based immunotherapy can be widely adopted. For instance, the non-specific targeting of the CAR T cells against normal tissues (on-target off-tumor toxicities) can be life-threatening: off-tumor toxicities against the lung, the brain, and the heart have caused multiple cases of deaths (*7–10*). Immunosuppressive corticosteroid therapy and suicide gene engineering are relatively effective in suppressing off-tumor toxicities and related cytokine release syndrome (CRS), but they fail to discriminate between beneficial T cell functions and toxic side effects (*11–13*). Synthetic biology and genetic circuits have been used to enhance specificity and reduce off-tumor toxicity by creating chemically inducible dimerization of split CARs, inhibitory CARs (iCARs), and SynNotch to control CAR activation (*8, 14–18*). However, given the extensive overlaps of antigens between solid tumors and normal tissues, especially those under conditions of tissue injury/inflammation (*19*), it remains very difficult to identify ideal antigens and their combinations to differentiate tumors from normal tissues. There is hence an urgent need for a high-precision control of CAR-T cells to confine the activation at local sites of solid tumors. Recently, we demonstrated that ultrasound signals can be amplified by microbubbles coupled to cells engineered with the mechanosensor Piezo1 to precisely control CAR T cell activations (*20*). However, the presence of microbubbles as cofactors limits the application of this system *in vivo*. Here, we have engineered a new class of inducible CAR T cells that can be remotely and directly controlled by FUS without any exogenous cofactor. We show that short-pulsed FUS stimulation can activate the engineered T cells at the desired time and location to suppress tumor growth *in vivo*.

## Results

### Heat-induced reporter gene activation

We propose to genetically engineer T cells with inducible CAR cassettes that can be remotely and directly activated, without any exogenous cofactor, by MRI-guided FUS at local tumor sites for recognizing and eradicating the tumor cells (Fig. 1a).

**Fig. 1.**
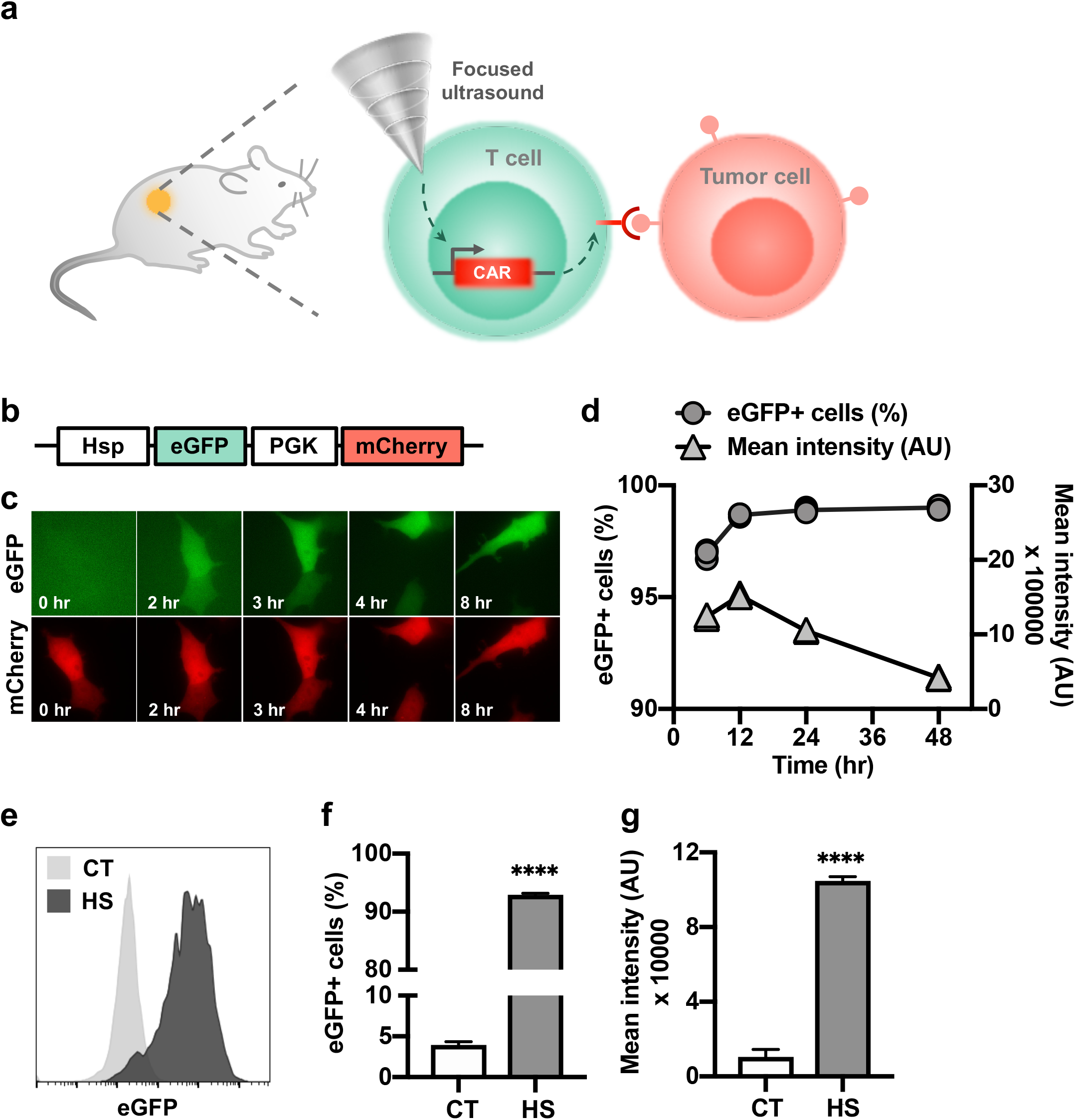
Heat-inducible gene activation. **(a)** Design of the FUS-controllable CAR T therapy technology. T cells engineered with the heat-inducible CAR and localized at the tumor region are activated by MRI-guided FUS for recognizing and eradicating target tumor cells. **(b)** Schematics of the dual-promoter eGFP reporter. **(c-d) (c)** Fluorescent images of inducible eGFP and constitutive mCherry, and **(d)** the percentage of eGFP+ cells and their mean fluorescence intensity after a 15-min HS at 43°C in HEK 293T cells containing the dual-promoter reporter. **(e-g)** Gene induction in primary human T cells with the dual-promoter eGFP reporter. **(e)** Representative flow cytometry profiles of eGFP expression. **(f)** The percentage of eGFP+ cells and **(g)** their mean fluorescence intensity. In (e-g), CT: without HS; HS: with a continuous 15-min HS. MCherry+ cells were gated for eGFP analysis. N = 3 repeats; error bar: SEM. ****: p < 0.0001.

We first tested the inducible activation of a reporter eGFP under the control of the heat-shock-protein promoter (Hsp). We assembled a dual-promoter reporter construct containing the Hsp-driven eGFP and a constitutive PGK-driven mCherry (Fig. 1b). HEK 293T cells infected with the reporter lentivirus (fig. S1a) were heated at 43°C for 15 min. Real-time fluorescence imaging revealed that the heat-induced eGFP expression started as early as 2 hr after heat shock (HS) and persisted throughout the course of observation (Fig. 1c and Movie S1). Quantitative tracking of the dynamics of heat-induced eGFP expression by flow cytometry showed that 97% of the cells expressed eGFP at 6 hr post HS, and the percentage increased to 99% at 12 hr and remained stable for 2 days, while the mean fluorescence intensity peaked at 12 hr followed by a steady decrease (Fig. 1d). We then investigated the inducible effect of HS in primary human T cells hosting the dual-promoter eGFP reporter (fig. S1b). A 15-min HS induced a strong eGFP expression in 92.9% of the engineered T cells, in contrast to a background of 3.9% in control cells without HS (Fig. 1, e and f). The mean fluorescence intensity of the eGFP+ cells in the HS group was 10-fold of that in the control group without HS (Fig. 1g).

### Heat-induced CAR expression and its functionality in Jurkat and primary human T cells

In order to convert the transient heat stimulation to a sustained gene activation and cellular functions for therapeutic actions, we integrated the Cre-lox gene switch into the inducible system. The design is composed of two constructs, one containing the Hsp-driven Cre recombinase and the PGK-driven membrane c-Myc tag for cell sorting (“inducible Cre”, Fig. 2a), and the other containing a lox-flanked “ZsGreen-STOP” sequence between a PGK promoter and an anti-CD19 CAR (“lox-stop CAR reporter”, Fig. 2a). As such, the excision of the “STOP” cassette mediated by the transient heat-induced Cre can cause a switch from ZsGreen to sustained CD19CAR production.

**Fig. 2.**
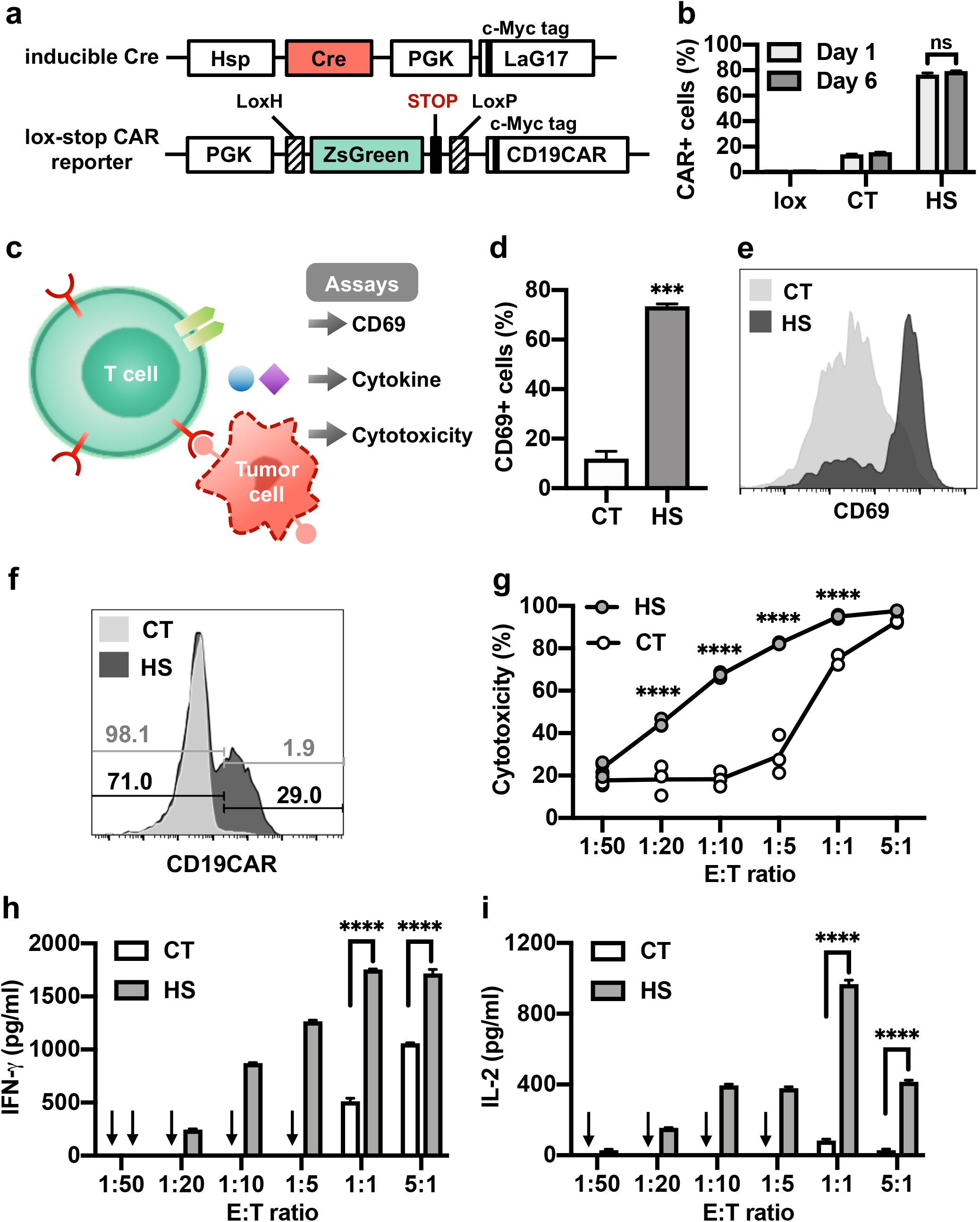
Heat-inducible CD19CAR expression and functional outcomes in Jurkat and primary T cells. **(a)** Schematics of the transgenes: the inducible Cre and the lox-stop CAR reporter. **(b)** Inducible CAR expression in Jurkat cells hosting the lox-stop CAR reporter alone (lox), or both transgenes in (a) with (HS) or without HS (CT). **(c)** Schematics of assays accessing the functionality of the heat-induced CAR T cells, including CD69 expression, cytotoxicity, and cytokine release. **(d)** The percentage of CD69+ cells in Jurkat with both transgenes in (a). **(e)** Representative flow cytometry data showing the histogram of CD69 expression in (d). **(f)** Representative histograms showing the percentage of CAR+ cells in primary T cells with both transgenes in (a). **(g)** The cytotoxicity of the T cells in (f) against Nalm-6 tumor cells at various E:T ratios. **(h-i)** Quantification of **(h)** IFN-γ and **(i)** IL-2 cytokine release associated with (g). Arrow: cytokine level not detectable. In (b) and (d) to (f), CT: without HS; HS: with a continuous 15-min HS. N = 3; error bar: SEM. ***: p < 0.001; ****: p < 0.0001; ns: no significant difference.

We first tested this heat-inducible gene switch system in Jurkat T cell lines (fig. S2a). A 15-min HS induced CAR expression in 76.6% of the cells when measured 24 hr after HS (Day 1), in contrast to a basal value of 14.0% in control cells without HS and a minimal leakage of 0.6% in cells infected with the lox-stop CAR reporter alone (Fig. 2b). The heat-induced CAR expression remained stable when measured 6 days after HS (Day 6, Fig. 2b). We further examined the functionality of the induced CD19CAR in engineered cells (Fig. 2c). Engineered Jurkat cells with (HS) or without (CT, control) a 15-min HS were co-cultured with CD19-expressing Nalm-6 tumor cells for 24 hr. Quantification of the expression level of CD69 (an early T cell activation marker) revealed a 73.4% CD69+ cell population in the engineered Jurkat cells in the HS group, in contrast to a 11.9% in the control group (Fig. 2, d and e). These results indicate that the HS-induced CD19CAR is efficient for functional changes in engineered Jurkat T cells.

We then examined our system in primary human T cells (Fig. 2, a and c; fig. S2b). CAR antibody staining showed that a 15-min HS induced CAR expression in 29% of the T cells, in contrast to 1.9% in control cells without HS (Fig. 2f). The heat-inducible CAR T cells were then co-cultured with firefly luciferase (Fluc)-expressing Nalm-6 cells at different effector-to-target (E:T) ratios for cytotoxicity assays. The luminescence of the remaining Nalm-6 cells was quantified after 24-hr co-culture. The heat-stimulated T cells (HS) demonstrated increased cytotoxicity with increased E:T ratio, with the largest difference in cytotoxicity between the HS and control (CT) T cells observed at E:T = 1:5, eliminating 82.9% and 29.3% of the target tumor cells, respectively (Fig. 2g). The heat-stimulated CAR T cells also released significantly higher concentrations of cytokines (IFN-γ and IL-2) than the control cells when co-cultured with Nalm-6 cells (Fig. 2, h and i), verifying the functional capability gained with the HS-induced CAR T cells.

While the continuous 15 min HS could lead to strong gene inductions (Figs. 1 and 2), it may cause toxicity to cells (*21*). We hence investigated the effect of different HS patterns in primary human T cells (fig. S3). Our results showed that longer HS resulted in more cell death; however, pulsed HS was able to alleviate this toxicity while achieving induction levels comparable to that in response to continuous HS with the same total heating time (fig. S3). In particular, a pulsed HS with 50% duty cycle and a total heating time of 15 min (fig. S3a, Pattern 2) caused a strong induction of eGFP expression in 91.4% of the engineered T cells, with a minimal toxicity as evidenced by the 92.2% cell viability measured 24 hr after HS (fig. S3, b to d). Therefore, we applied this HS pattern (fig. S3a, Pattern 2) for *in vivo* therapeutic studies.

### MRI-guided FUS-induced gene activation in phantom and *in vivo*

MRI-guided FUS enables the delivery of thermal energy *in vivo* at confined local regions with high spatiotemporal resolutions (*3, 4*). We integrated an MRI-guided FUS system (Image Guided Therapy) with a 7T MRI as described in Methods. An annular array transducer is placed above the target region of the object to be heated (phantoms or small animals) in the MRI bore. MR images are acquired and transferred to Thermoguide software to calculate the temperature of the target region in real-time, which is fed back to the PID controller to automatically regulate the output power of the FUS generator, maintaining the temperature of the target region at the desired level (Fig. 3a, fig. S4).

**Fig. 3.**
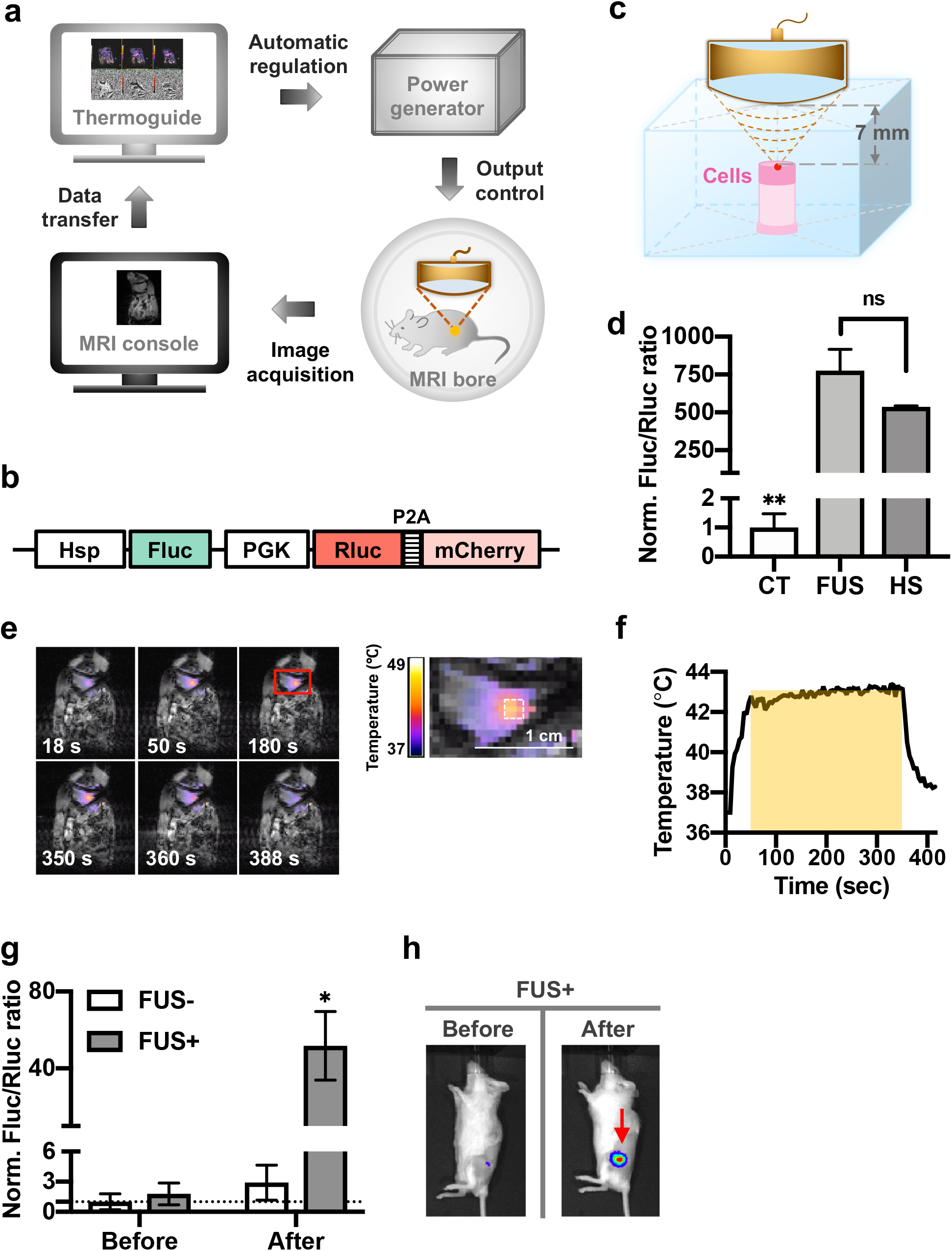
MRI-guided FUS-inducible gene activation in phantom and *in vivo*. **(a)** Schematics of the MRI-guided FUS system. **(b)** The dual-luciferase reporter containing the inducible Hsp-driven Fluc and constitutive PGK-driven Rluc fused with mCherry. **(c)** The experimental setup of FUS stimulation on cells in a tofu phantom. **(d)** Gene induction level in Nalm-6 cells containing the dual-luciferase reporter with three pulses of 5-min heating by MRI-guided FUS in tofu phantom (FUS) or by thermal cycler (HS). CT: without heating. Gene induction level is quantified by the Fluc/Rluc ratio and normalized to CT. N = 3. **(e)** Left: color-coded temperature map superimposed on MRI images at different time points during a 5-min FUS stimulation at 43°C on the hindlimb of an anesthetized mouse. Right: close-up of the red rectangle region on the left. The dotted white square outlines the region of interest (ROI) for temperature regulation. The average temperature of the ROI during FUS stimulation in (e). The yellow shadow represents the predefined target temperature (43°C) and duration (300 sec) of FUS stimulation. Gene induction *in vivo* by MRI-guided FUS. Nalm-6 cells containing the dual-luciferase reporter were injected subcutaneously into NSG mice followed by FUS stimulation. FUS+ or FUS−: with or without two pulses of 5-min FUS stimulation at 43°C. Gene induction was quantified by the *in vivo* Fluc/Rluc ratio and normalized to the “FUS−, before” group, as indicated by the dotted line. N = 4 mice. **(h)** Representative bioluminescence images of Fluc expression before and after FUS stimulation in (g). Error bar: SEM. *: p < 0.05; **: p < 0.01; ns: no significant difference.

We transduced Nalm-6 cells with a lentiviral dual-luciferase reporter containing inducible Fluc and constitutive Rluc (Hsp-Fluc-PGK-Rluc-mCherry; Rluc, *Renilla* luciferase; Fig. 3b) and embedded them in a tofu phantom approximately 7 mm deep from the top surface (Fig. 3c and Methods). We then focused the ultrasound on the embedded cells by changing the focal distance in the z direction. Three pulses of 5-min FUS stimulations caused a significant induction of gene expression as quantified by the Fluc/Rluc ratio of the cells assayed 8 hr later (Fig. 3d, Methods). The induction level is comparable to that of the positive control using thermal cycler with the same heating pattern (Fig. 3d), suggesting the acoustogenetic approach can remotely control gene activation in engineered cells with high efficiency.

We then used MRI-guided FUS to control local temperature *in vivo* in mouse (Fig. 3, e and f and Movie S2) and tested the FUS-induced gene activation using Nalm-6 cells with the dual-luciferase reporter *in vivo*. Significant gene induction was observed in the implanted cells with only two pulses of 5-min FUS stimulation (FUS+, after), in comparison to the basal level (FUS+, before) and the control groups (FUS−, before and after) (Fig. 3, g and h).

### FUS-inducible tumor cytotoxicity of the engineered CAR T cells *in vivo*

We next tested the tumor cytotoxicity of the FUS-inducible CAR T cells *in vivo*. We subcutaneously injected Nalm-6 cells (Fluc+) on both hindlimbs of NSG mice to generate matched bilateral tumors (Fig. 4a). Four days later, engineered CD19CAR T cells were subcutaneously injected at both tumor sites locally, followed by three pulses of 5-min FUS stimulation at 43°C on the left but not on the right tumor (Fig. 4a). Bioluminescence imaging revealed that FUS significantly suppressed tumor growth (Fig. 4, b and c). The results on the two tumors (FUS+ and FUS−) on the same mouse indicate that the FUS-activated CAR T cells at the local site had negligible off-site effects in attacking the distal tissues on the contralateral hindlimb expressing the same antigens. We further performed a control experiment subjecting mice carrying bilateral tumors to FUS stimulation on one side, with neither site subjected to the injection of engineered CAR T cells (fig. S5a). The tumors with or without FUS stimulation exhibited similar growth profiles, indicating that FUS itself (with the chosen pattern) had no impact on tumor growth (fig. S5, b and c). Therefore, our results demonstrated that FUS can be used to precisely control the cytotoxicity of the engineered CD19CAR T cells *in vivo* against target tumor cells.

**Fig. 4.**
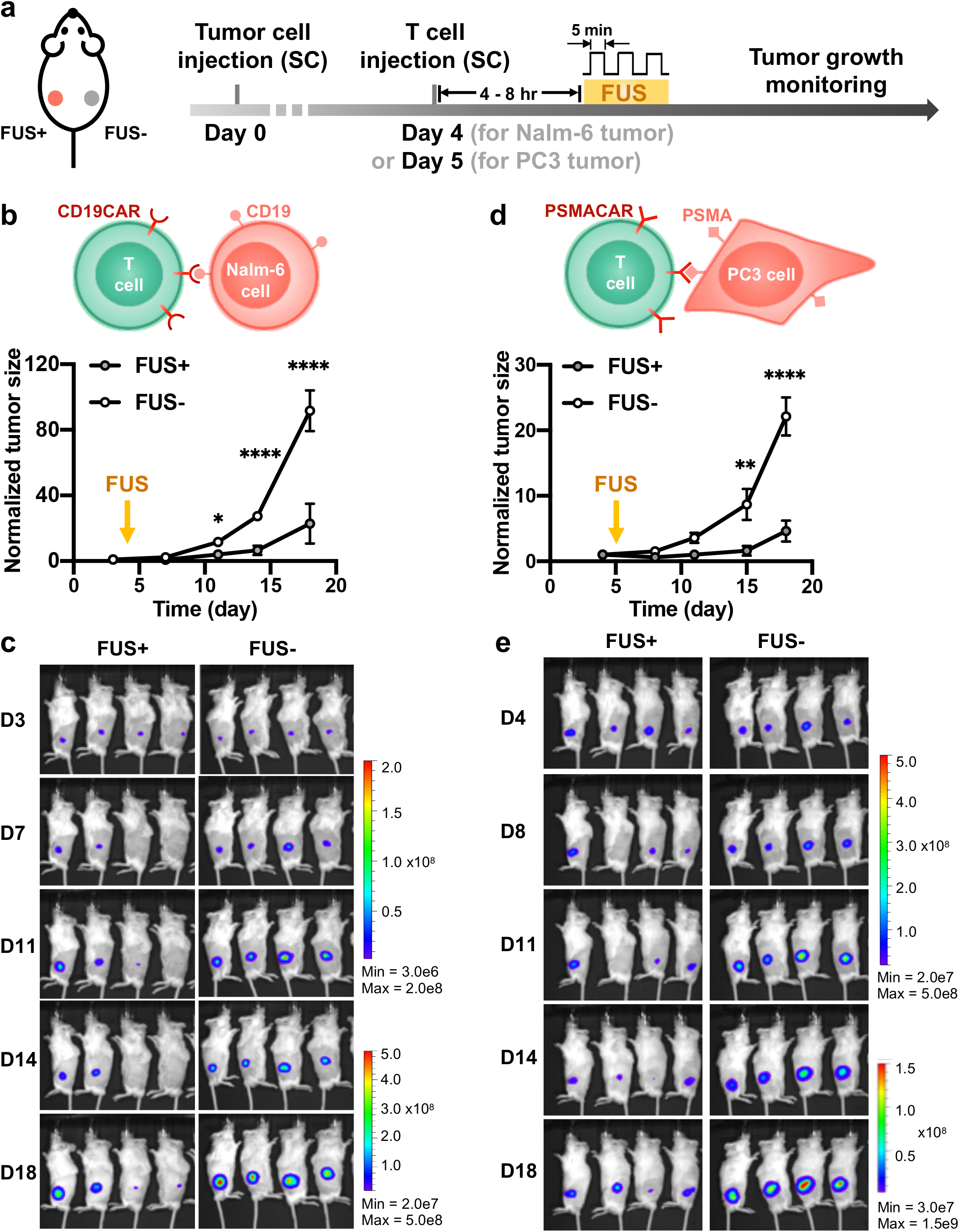
FUS-controllable tumor suppression by the engineered CAR T cells *in vivo*. **(a)** Timeline of the *in vivo* experiment using NSG mouse bearing matched bilateral tumors as the animal model. The tumor on the left flank received FUS stimulation (FUS+) and the one on the right received no FUS (FUS−) following injection of engineered CAR T cells. **(b-e)** The quantified tumor growth and representative bioluminescence images of **(b-c)** Nalm-6 tumors or **(d-e)** PC3 tumors with (FUS+) or without (FUS−) FUS stimulation. Tumor size was quantified using the integrated Fluc luminescence intensity of the tumor region and normalized to that of the same tumor on the first measurement. N = 4 mice. Error bar: SEM. *: p < 0.05; **: p < 0.01; ****: p < 0.0001.

We further tested this acoustogenetic technology in controlling inducible CAR T cells against other types of tumors, particularly solid tumors. We engineered solid tumor human prostate cancer PC3 cells to express the prostate-specific membrane antigen (PSMA) and Fluc, and engineered primary human T cells with the Cre-lox mediated heat-inducible anti-PSMA CAR (PSMACAR; fig. S6a). We verified the functionality of the heat-inducible PSMACAR T cells through *in vitro* co-culture cytotoxicity assays and the associated cytokine assays (fig. S6, b to d). We then generated matched bilateral subcutaneous PC3 tumors (PSMA+, Fluc+) in NSG mice; five days later we subcutaneously injected heat-inducible PSMACAR T cells next to the tumor sites on both sides. The tumor regions on the left side were treated with three pulses of 5-min FUS, while those on the right remained unstimulated. Consistently, the tumors with FUS stimulation showed significantly inhibited growth as compared to the controls (Fig. 4, d and e). We further harvested the tumor tissues and quantified the related mRNA amount. The CD3 mRNA in the FUS-treated tumors averaged 3-fold of that in the untreated ones, indicating more T cell infiltration in the FUS-treated solid prostate tumors (fig. S7a). Moreover, the amount of Cre-mediated recombined CAR mRNA in the FUS-treated tumors was 9-fold of that in the untreated controls, verifying the FUS-induced DNA recombination and subsequent CAR expression in the engineered T cells at the tumor sites (fig. S7, b and c, and Methods). These results demonstrated the efficacy of FUS-based acoustogenetics in remote control of CAR T cells for treating different types of tumors *in vivo*, including solid tumors of prostate cancer.

## Discussion

We developed an FUS-based acoustogenetic approach to remotely control, without any exogenous cofactor, the genetically engineered T cells capable of perceiving ultrasound signals and transducing them into genetic and cellular activations for therapeutic applications *in vivo*. This acoustogenetics technology enables the activation of CAR T cells at confined tissue regions, thus allowing the targeting of the less ideal antigens without causing non-specific off-site cytotoxicity. This is of critical importance given the extensive overlap of antigens between tumors and normal cells, particularly those under conditions of tissue injury and inflammation. The short-pulsed patterns of FUS stimulation should also minimize potential detrimental effects of hyperthermia and induce transient expression of synthetic protein regulators to circumvent severe immune responses. This acoustogenetic approach is highly modular, with the target CAR genes switchable to aim at different cancer types.

We employ the Cre-mediated gene switch to convert transient FUS inputs into sustained outputs of genetic and cellular activities for sufficient therapeutic efficiency. The nature of local activation should limit the number of activated cells off the tumor site and the potential non-specific cytotoxicity against normal tissues, as evidenced in our results (Fig. 4, b to e; if the Cre-mediated permanent activation of CAR becomes an issue in the future, degradation domains such as dihydrofolate reductase (DHFR) can be fused to CAR to control the protein lifetime with an FDA-approved drug methotrexate (*22*). This “AND” gate with FUS and methotrexate should enhance the precision of controllable CAR T immunotherapy.

We anticipate that comparable therapeutic outcomes can be achieved in a reversible heat-inducible system without the Cre-lox gene switch, but this may require multiple rounds of FUS stimulation. In such a system, Hsp directly drives the production of CAR (Hsp-CAR) under FUS stimulation. Upon the withdrawal of FUS stimulation, HSFs gradually dissociate from Hsp, returning Hsp and its downstream transcriptional activities to the resting state. This recovery process is relatively fast, within 45 min after HS for *Drosophila* Hsp70 and approximately 60 min after HS for human Hsp70 (*23, 24*). The dynamics of this heat-induced CAR expression hence largely depends on its protein lifetime, with the half-life of GFP-tagged CAR reported to be around 8 hr (*16*). Therefore, repeated FUS stimulation can be applied to maintain the CAR expression in the T cells (and hence their cytotoxicity) for a sustained period of time. We tested this concept by applying a 10-min HS every 48 hr in T cells with Hsp-eGFP, and indeed observed oscillatory patterns of the induced eGFP expression (fig. S8). We anticipate that T cells with a simple Hsp-CAR can also be repeatedly activated by FUS to achieve sustained CAR expression and cytotoxicity for a desired period of time or until tumor elimination. Such a reversible FUS-inducible system can further prevent “on-target off-tumor” toxicity of canonical CAR T therapy, as the T cells leaving the tumor site will no longer receive FUS stimulation and gradually lose CAR molecules. The tunable FUS pulses should also allow the precise control of the temporal activation patterns of CAR T cells for an optimized killing efficiency with controllable exhaustion.

We chose local injection at the tumor site to deliver T cells *in vivo*. Local administration of CAR T cells has been tested in animals and patients to overcome the obstacle of T cell homing associated with intravenous delivery, and has achieved promising therapeutic effects (*16, 25, 26*). For example, since the prostate is positioned near critical organ structures including urethra and neurovascular bundles, surgery or radiation therapy targeting the whole prostate gland to treat the prevalent locally-progressed prostate cancer (*27*) may cause adverse effects that would significantly impact quality of life (*28, 29*). Local delivery and activation of inducible CAR T cells using clinically available MRI-guided FUS systems should allow, without any exogenously added nanoparticle or cofactor, a high degree of precision and safety in eradicating tumor cells in these patients harboring locally progressed prostate cancer (*29*). In cases where intravenous delivery is required, it is also possible to equip the FUS-inducible CAR T cells with additional antigen binders and/or chemokine receptors to promote trafficking, infiltration, and the enrichment of these engineered cells at the tumor site before FUS activation (*30, 31*).

The short-pulsed stimulation and the biocompatible Hsp capable of inducing transient expressions of different synthetic protein regulators can potentially enhance the safety of gene therapy, circumventing detrimental host immune response. For instance, CRISPR-Cas9 proteins have been a powerful tool for research in genetic and epigenetic engineering, but can evoke adaptive immune responses and tissue damage *in vivo*, and are therefore potentially pathogenic if applied to correct inherited genetic defects to treat diseases (*32*). Protein engineering to remove immunogenic epitopes and humanize these synthetic proteins to circumvent this issue can be difficult owing to the high diversity of the human leukocyte antigen (HLA) loci (*33*). Using our acoustogenetic approach, the transiently induced Hsp-driven synthetic regulators (e.g. Cas9) can be cleared in a timely manner to mitigate or evade the adaptive immune response, hence offering a new option for gene therapy.

Each component of this FUS-based acoustogenetics, i.e. ultrasound devices, molecular thermo-sensors, and genetic/epigenetic transducing modules, is highly modular and will continue to evolve for greater precision and reduced immunogenicity. In fact, stretchable electronic circuits are being developed to fabricate wearable patches of ultrasound transducers (*34*). The leverage of technological advancements of different fields into FUS-based acoustogenetics should be able to drive the development of these fields to open up new frontiers. We envision that the current state of acoustogenetics is analogous to optogenetics at its infancy. Before the functional demonstration of channelrhodopsin in neuronal cells (*35*), it was challenging to manipulate molecular activities in live cells at high spatiotemporal resolutions. With the technological integration and convergence of optics, genetic circuits, and light-sensitive proteins, optogenetics is rapidly reaching its full potential. Based on this analogy, acoustogenetics may undergo a similar trajectory to provide a broadly applicable method and usher in an era of applying ultrasound for the direct, remote, and noninvasive control of genetically engineered cells for therapeutics.

## Methods

### Cloning

Plasmids used in this paper are listed in Table S1. Cloning strategies include Gibson Assembly (NEB, E2611L) and T4 ligation (NEB, M0202L). PCR was performed using synthesized primers (Integrated DNA Technologies) and Q5 DNA polymerase (NEB, M0491). The sequences of the constructed plasmids were verified by Sanger sequencing (Genewiz).

### General cell culture

HEK 293T cells were cultured in DEME (Gibco, 11995115) with 10% FBS (Gibco, 10438026) and 1% Penicillin-Streptomycin (Gibco, 15140122). Jurkat, Nalm-6, and PC3 cells were cultured in RPMI 1640 (Gibco, 22400105) with 10% FBS and 1% P/S. Primary human T cells were cultured in complete RPMI 1640 supplemented with 100 U/mL recombinant human IL-2 (PeproTech, 200-02). Cells were cultured at 37°C in a humidified 5% CO_2_ incubator.

### Staining and flow cytometry

Staining of cell surface markers (e.g., c-Myc, CD69, etc) for flow cytometry was performed using fluorophore-conjugated antibodies according to manufacturers’ protocols. In general, cells were washed twice and resuspended in 100 µL wash buffer (PBS + 0.5% BSA) containing the suggested amounts of antibodies, incubated in dark at room temperature for suggested durations, and washed three times before being analyzed using a BD flow cytometer. Gating was based on non-engineered cells with the same staining. Flow cytometry data were analyzed using FlowJo software (Tree Star).

### *In vitro* heat shock

For Fig. 1c and Movie S1, cells seeded in a glass bottom dish were heated at 43°C for 15 min using a heating stage (Instec) integrated with a Nikon Eclipse Ti inverted microscope. Images were acquired in real-time to obtain the kinetics of the induced fluorescent protein. For the remainder of the *in vitro* heat shock (HS) experiment, unless otherwise specified, cells were washed and resuspended in cell culture medium at a concentration of 2 × 10^6^ cells/mL, aliquoted into 8-strip PCR tubes with 50 µL/tube, and heat shocked at 43°C using a thermal cycler (Bio-Rad, 1851148) with various patterns as indicated (Table S2). Cells were returned to standard culture condition after HS. The gene induction levels were quantified by flow cytometry 12 hr after HS in Fig. 1, f and g, and fig. S3, d and e.

### Engineered cells

The engineered cells (excluding primary human T cells) used in this work are listed in Table S3. Lentiviruses were used to deliver engineered genes into the cells. Fluorescence-activated cell sorting (FACS), when needed, was performed at UCSD Human Embryonic Stem Cell Core Facility by professional technicians following standard protocols.

### Quantification of CAR expression in Jurkat cells

Jurkat cells were either transduced with a lentiviral cocktail (inducible Cre and lox-stop CAR reporter, Fig. 2a) followed by indicated HS (Fig. 2b), or transduced with the lox-stop CAR reporter lentivirus alone without HS. CAR expression was quantified by CAR antibody staining (an anti-mouse IgG, F(ab’)_2_ fragment specific antibody; Jackson ImmunoResearch, 115-606-072) and flow cytometry 24 hr after HS. Live single cells were gated for CAR expression analysis. Non-engineered Jurkat cells were stained with the same antibody to generate the CAR+ gate.

### Quantification of CD69 expression in Jurkat cells

Jurkat cells transduced with a lentiviral cocktail (inducible Cre and lox-stop CAR reporter, Fig. 2a) were treated with or without HS at 43°C for 15 min, and co-cultured with target tumor cells for 24 hr. The cells were then stained by an APC anti-human CD69 antibody (BioLegend, 310910) and analyzed by flow cytometry. ZsGreen+ cells (representing the engineered Jurkat cells) were gated for analysis of CD69 expression. Non-engineered Jurkat cells co-cultured with target tumor cells were stained with the same antibody to generate the CD69+ (APC+) gate.

### Isolation, culture, transduction and MACS of primary human T cells

Human peripheral blood mononuclear cells (PBMCs) were isolated from buffy coats (San Diego Blood Bank) using Lymphocyte Separation Medium (Corning, 25-072-CV) following the manufacturer’s instructions. Primary human T cells were isolated from PBMCs using Pan T Cell Isolation Kit (Miltenyi, 130-096-535) and activated with Dynabeads® Human T-Expander CD3/CD28 (Gibco, 11141D). Three days later, lentivirus concentrated using PEG-it (SBI, LV825A-1) was added to the T cells at MOI = 10, followed by spinoculation in a 96-well plate coated with Retronectin (Takara, T100B). T cells were further expanded and dynabeads were removed prior to downstream procedures (e.g., magnetic-activated cell sorting (MACS), *in vitro* HS, *in vivo* injection, etc.).

For *in vitro* and *in vivo* cytotoxicity studies, T cells were transduced with a lentiviral cocktail of inducible Cre and lox-stop CAR reporter (Fig. 2a). MACS was performed using Anti-c-Myc-Biotin antibodies and Anti-Biotin microbeads (Miltenyi, 130-092-471 and 130-097-046) following the manufacturer’s instructions to enrich c-Myc+ cells. A representative double positive efficiency after MACS is 69%, 95% for the c-Myc+ and 71.4% for the ZsGreen+ cells (fig. S2b). CAR expression in the engineered inducible CAR T cells with or without HS was quantified using the CAR antibody as described above.

### Luciferase-based cytotoxicity assay

A constant number of 5 × 10^4^ Fluc+ Nalm-6 cells were mixed with engineered primary human T cells with or without HS (pre-washed and resuspended with complete RPMI without IL-2) at effector-to-target (E:T) ratios of 1:50, 1:20, 1:10, 1:5, 1:1, 5:1 or no T cells (“target cell only”). The mixtures were then cultured in round bottom 96 well plates for 24 hr, centrifuged to remove the supernatant (which was harvested for quantification of cytokine production), and assayed with the Bright-Glo™ Luciferase Assay System (Promega, E2610) following the manufacturer’s instructions to quantify the luminescence of each sample. The cytotoxicity (%) of Sample X was calculated as (1−Luminescence of X / Luminescence of “target cell only”) × 100%.

For cytotoxicity assay using PC3 cells as the target, 1 × 10^4^ PSMA+ Fluc+ PC3 cells were seeded onto TC-treated flat bottom 96 well plates (Corning, 3603). Except for “target cell only” wells, engineered primary human T cells with or without HS (washed and resuspended with complete RPMI without IL-2) were added 6 hr later at E:T ratios of 1:10, 1:5, 1:1, 5:1, 10:1, 20:1. The luminescence was quantified 24 hr after co-culture as described above.

### Quantification of cytokine production

The supernatant of effector-target cell co-culture was harvested. The concentrations of cytokines IL-2 and IFN-γ were quantified using the corresponding ELISA kits (BD, 555190 and 555142).

### T cell viability assay

Non-engineered primary human T cells were heat shocked as described above and kept under normal culture condition for 24 hr. Cell viability was then assessed using the FITC Annexin V Apoptosis Detection Kit I (BD, 556547) following the manufacturer’s instructions. The cells stained negative for both Annexin V and PI were counted as live cells.

### MRI-guided FUS

The MRI-guided FUS system is composed of a 1.5 MHz 8-element annular array transducer, a 16-channel broadband RF generator, a piezo motor-based X-Y positioning stage, and a degassing and water circulation system (Image Guided Therapy, France). MR images acquired using a Bruker 7T MRI system were transferred to ThermoGuide software (Image Guided Therapy, France) to generate phase images and real-time temperature maps. Using PID controller, the software automatically regulates the output power of the generator to maintain the temperature at the focal spot at a desired value as described elsewhere (*6, 36*).

Animal experiments were performed following Protocol S15285 approved by UCSD IACUC. NSG mice (6-8 weeks old) were purchased from Jackson Laboratory (JAX) and shaved prior to FUS stimulation. Anesthesia was induced using 2% isoflurane-oxygen mixture and maintained with 1.5% isoflurane-oxygen mixture during FUS stimulation. The mouse was laid on its side on an MR bed containing an agarose gel pad and a surface coil. A pressure pad was placed under the mouse to monitor its respiration rate, and a rectal thermal probe was used to provide feedback for the delivering of warm air into the bore to maintain the mouse’s core temperature at approximately 37°C. The ultrasound transducer was positioned right above the targeted region on the mouse’s hindlimb. Thin layers of SCAN ultrasound gel (Parker labs) were applied at the skin-transducer and skin-bed interfaces.

The ThermoGuide software regulates the temperature in a 3 × 3 pixel square (3 - 4 mm^2^) centered at the ultrasound focus (Fig. 3e). A PID controller is used to maintain the average temperature of the target square at 6°C above reference by controlling the output power of the FUS generator, with the reference temperature being 37°C as measured by the rectal thermal probe. As such, the MRI-guided FUS enabled temperature elevation to 43°C locally at the focal area in the hindlimb of an anesthetized mouse.

### FUS stimulation in tofu phantom

For FUS stimulation on cells in the tofu phantom, Nalm-6 cells were lentivirally transduced with the dual-luciferase reporter (Fig. 3b, Hsp-Fluc-PGK-Rluc-mCherry) and FACS-sorted. The cells were resuspended in culture medium and mixed with matrigel (Corning, 354262) at 1:1 volume ratio on ice. Extra-firm tofu was cut into a 15-mm thick pad, and an 8-mm deep hole of 8-mm diameter was drilled from the top. A microcentrifuge tube of 7.5-mm diameter (Fisherbrand, 05-408-120) was cut to 8-mm long by removing the lid and the conical bottom, and was inserted into the hole in the tofu phantom. Cell-matrigel mixture of 150 µL was added into the hole (~3 mm thick) and allowed to gel at room temperature. The rest of the hole and the gap between the tube and the tofu phantom were filled up with matrigel. After gelation, the assembly was inverted and positioned onto the MR bed containing the surface coil. The ultrasound transducer was positioned above the tofu phantom with its center aligned with that of the tube. Thin layers of ultrasound gel were applied at the tofu-transducer and tofu-bed interfaces. A thermal probe was inserted into the distal end of the tofu phantom to provide reference temperature readings.

MR images of the assembly were acquired and transferred to ThermoGuide to calculate the theoretical ultrasound focal position. Test FUS shots were delivered to determine the actual focal position. Steering was applied to focus the ultrasound at the region immediately above the cells. Three pulses of 5-min FUS stimulations at 43°C were applied. The cell-matrigel mixture was then recovered from the tube, placed in cell culture medium, and returned to a standard 37°C cell culture incubator. After 6 hr, the culture was centrifuged to remove the supernatant, and the cell-matrigel pellet was incubated in a Cell Recovery Solution (Corning, 354253) at 4°C for 1 hr to retrieve the Nalm-6 cells from matrigel. The Fluc and Rluc luminescence of the cells was quantified using the Dual-Luciferase® Reporter Assay System (Promega, E1910) following the manufacturer’s instructions.

### *In vivo* bioluminescence imaging

*In vivo* bioluminescence imaging (BLI) was performed using an In vivo Imaging System (IVIS) Lumina LT Series III (PerkinElmer). For Fluc imaging, 150 mg/kg D-Luciferin (GoldBio, LUCK) was administered intraperitoneally (i.p.). BLI started 10 min after substrate injection until peak signal was acquired. For Rluc imaging, 200 µL 0.295 mM ViviRen™ (Promega, P1232) (*37*) was administered i.p. BLI started 15 min after substrate injection until peak signal was acquired. BLI of Fluc and Rluc in the same mouse, when needed, was performed 4 hr apart. Images were analyzed using Living Image software (PerkinElmer), and the integrated Fluc luminescence intensities within regions of interest were quantified to represent tumor sizes.

### FUS-inducible gene activation *in vivo*

NSG mice (male, 6-8 weeks old) were subcutaneously injected with 2 × 10^6^ dual-luciferase reporter Nalm-6 cells at the hindlimb. One week later, the experimental mice received two pulses of 5-min FUS stimulation at 43°C targeted at the implanted cells, while the control mice remained unstimulated. The *in vivo* Fluc and Rluc luminescence was quantified 4 hr before and 12 hr after FUS stimulation, as described above.

### *In vivo* tumor cytotoxicity of FUS-inducible CAR T cells

NSG mice (male, 6-8 weeks old) were subcutaneously injected with 2 × 10^5^ Fluc+ Nalm-6 cells (or 2 × 10^5^ PSMA+ Fluc+ PC3 cells, for PC3 tumors) on both hindlimbs to generate matched bilateral tumors. Four days later (or five days later, for PC3 tumors), 1 × 10^6^ inducible primary human CAR T cells prepared as described above were injected subcutaneously and locally at tumor regions. Within 4 - 8 hr after T cell injection, three pulses of 5-min FUS stimulation targeted at 43°C were applied on the left tumor region as described above, while the tumor on the right hindlimb received no FUS stimulation to serve as the control. Tumor aggressiveness was monitored by BLI twice a week as described above until euthanasia criteria were met.

### Quantification of mRNA expression in tumor tissue

PC3 tumors (Fig. 4, d and e) were harvested 22 days after tumor implantation (17 days after T cell injection and FUS stimulation). The tumors were disrupted and homogenized, and the same amount of lysate from each tumor was used to extract total RNA with the RNeasy Mini Kit (Qiagen, 74104) followed by reverse transcription using the same amount of template RNA. Quantitative PCR (qPCR) was performed using iTaq™ Universal SYBRRTM Green Supermix (Bio-Rad, 1725121), the same amount of template cDNA, and the specific primers described below. The mRNA levels were normalized to 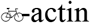.

The first pair of specific primers were designed on human CD3γ chain to detect the presence of human T cells. The second pair of specific primers were designed based on the lox-stop PSMACAR reporter sequence to reflect CAR expression after FUS-induced Cre recombination (fig. S6, a and b). The forward primer anneals from −60 bp of the mouse PGK promoter, downstream of the transcription starting site (TSS), and the reverse primer anneals from +20 bp of the PSMACAR gene (*38*). With the presence of FUS-induced Cre recombinase, the sequence from the second half of LoxH to the first half of LoxP will be excised, resulting in a 200 bp qPCR product. Without Cre-mediated recombination, this pair of primers will theoretically generate a 984-bp fragment. We adopted a two-step qPCR protocol with combined annealing/extension at 60°C for only 15 sec to ensure the specific amplification of the 200-bp fragment, but not the 984 bp fragment, as confirmed by gel electrophoresis and Sanger sequencing of the qPCR product (fig. S6, c and d; sequence alignment performed in Serial Cloner). Therefore, the second pair of specific primers can detect the successfully recombined CAR mRNA amount.

### Statistics

One-way ANOVA followed by Tukey’s multiple comparisons test is used for Figs. 1, f and g, and fig. S3, b and d to e. Student’s t-test is used for Fig. 2d. Two-way ANOVA followed by Sidak’s multiple comparisons test is used for Figs. 2, b, g to I, 3g, 4, b and d, fig. S4b, S5, b to d.

## Acknowledgments

We thank Dr. Franck Couillaud (University of Bordeaux, France) for providing the Hsp template and Dr. Michel Sadelain (Sloan Kettering Institute, USA) for the PSMA scFv and PSMA constructs and the Nalm-6 cells. We also thank Dr. Erik Dumont and Ms. Stéphanie Hoarau-Recco (Image Guided Therapy, France) for their most valuable help on the FUS system.

## Author contributions

Y.Wu, S.C. and Y.Wang designed research; Y.Wu, Y.L, Z.H., X.W., Z.J., J.L., P.L., L.Z., M.A., Y.P., R.B., A.J. performed research; Y.Wu and Y.L. analyzed data; Y.Wu, T. L., S.C. and Y.Wang wrote the manuscript. All authors reviewed the manuscript and have given approval to the final version of the manuscript.

## Data and materials availability

All data is available in the main text or the supplementary materials.

## References

1. R. Y. Tsien, Imagining imaging’s future. Nat Rev Mol Cell Biol Suppl, SS16–21 (2003).

2. M. Thanou, W. Gedroyc, MRI-Guided Focused Ultrasound as a New Method of Drug Delivery. J Drug Deliv 2013, 616197 (2013).

3. R. Deckers et al., Image-guided, noninvasive, spatiotemporal control of gene expression. Proceedings of the National Academy of Sciences of the United States of America 106, 1175–1180 (2009).

4. E. Guilhon et al., Image-guided control of transgene expression based on local hyperthermia. Mol Imaging 2, 11–17 (2003).

5. S. Wang, V. Zderic, V. Frenkel, Extracorporeal, low-energy focused ultrasound for noninvasive and nondestructive targeted hyperthermia. Future Oncol 6, 1497–1511 (2010).

6. D. I. Piraner, M. H. Abedi, B. A. Moser, A. Lee-Gosselin, M. G. Shapiro, Tunable thermal bioswitches for in vivo control of microbial therapeutics. Nature chemical biology 13, 75–80 (2017).

7. M. L. Davila et al., Efficacy and toxicity management of 19-28z CAR T cell therapy in B cell acute lymphoblastic leukemia. Science translational medicine 6, 224ra225 (2014).

8. D. Chakravarti, W. W. Wong, Synthetic biology in cell-based cancer immunotherapy. Trends in biotechnology 33, 449–461 (2015).

9. M. V. Maus, S. A. Grupp, D. L. Porter, C. H. June, Antibody-modified T cells: CARs take the front seat for hematologic malignancies. Blood 123, 2625–2635 (2014).

10. R. A. Morgan et al., Case report of a serious adverse event following the administration of T cells transduced with a chimeric antigen receptor recognizing ERBB2. Molecular therapy : the journal of the American Society of Gene Therapy 18, 843–851 (2010).

11. G. Akpek, S. M. Lee, V. Anders, G. B. Vogelsang, A high-dose pulse steroid regimen for controlling active chronic graft-versus-host disease. Biol Blood Marrow Transplant 7, 495–502 (2001).

12. A. Di Stasi et al., Inducible apoptosis as a safety switch for adoptive cell therapy. The New England journal of medicine 365, 1673–1683 (2011).

13. M. Themeli, M. Sadelain, Combinatorial Antigen Targeting: Ideal T-Cell Sensing and Anti-Tumor Response. Trends Mol Med 22, 271–273 (2016).

14. J. H. Cho, J. J. Collins, W. W. Wong, Universal Chimeric Antigen Receptors for Multiplexed and Logical Control of T Cell Responses. Cell 173, 1426–1438 e1411 (2018).

15. V. D. Fedorov, M. Themeli, M. Sadelain, PD-1- and CTLA-4-based inhibitory chimeric antigen receptors (iCARs) divert off-target immunotherapy responses. Science translational medicine 5, 215ra172 (2013).

16. K. T. Roybal et al., Precision Tumor Recognition by T Cells With Combinatorial Antigen-Sensing Circuits. Cell 164, 770–779 (2016).

17. C. Y. Wu, K. T. Roybal, E. M. Puchner, J. Onuffer, W. A. Lim, Remote control of therapeutic T cells through a small molecule-gated chimeric receptor. Science 350, aab4077 (2015).

18. M. M. D’Aloia, I. G. Zizzari, B. Sacchetti, L. Pierelli, M. Alimandi, CAR-T cells: the long and winding road to solid tumors. Cell Death Dis 9, 282 (2018).

19. S. I. Grivennikov, F. R. Greten, M. Karin, Immunity, inflammation, and cancer. Cell 140, 883–899 (2010).

20. Y. Pan et al., Mechanogenetics for the remote and noninvasive control of cancer immunotherapy. Proceedings of the National Academy of Sciences of the United States of America 115, 992–997 (2018).

21. I. C. Miller, M. Gamboa Castro, J. Maenza, J. P. Weis, G. A. Kwong, Remote Control of Mammalian Cells with Heat-Triggered Gene Switches and Photothermal Pulse Trains. ACS Synth Biol 7, 1167–1173 (2018).

22. M. V. Raimondi et al., DHFR Inhibitors: Reading the Past for Discovering Novel Anticancer Agents. Molecules 24, (2019).

23. K. Abravaya, B. Phillips, R. I. Morimoto, Attenuation of the heat shock response in HeLa cells is mediated by the release of bound heat shock transcription factor and is modulated by changes in growth and in heat shock temperatures. Genes Dev 5, 2117–2127 (1991).

24. S. K. Ghosh, A. Missra, D. S. Gilmour, Negative elongation factor accelerates the rate at which heat shock genes are shut off by facilitating dissociation of heat shock factor. Mol Cell Biol 31, 4232–4243 (2011).

25. M. Martinez, E. K. Moon, CAR T Cells for Solid Tumors: New Strategies for Finding, Infiltrating, and Surviving in the Tumor Microenvironment. Front Immunol 10, 128 (2019).

26. P. Sridhar, F. Petrocca, Regional Delivery of Chimeric Antigen Receptor (CAR) T-Cells for Cancer Therapy. Cancers (Basel) 9, (2017).

27. U. Mahmood et al., Current clinical presentation and treatment of localized prostate cancer in the United States. J Urol 192, 1650–1656 (2014).

28. H. B. Musunuru et al., Active Surveillance for Intermediate Risk Prostate Cancer: Survival Outcomes in the Sunnybrook Experience. J Urol 196, 1651–1658 (2016).

29. A. R. Rastinehad et al., Gold nanoshell-localized photothermal ablation of prostate tumors in a clinical pilot device study. Proc Natl Acad Sci U S A 116, 18590–18596 (2019).

30. M. Boice et al., Loss of the HVEM Tumor Suppressor in Lymphoma and Restoration by Modified CAR-T Cells. Cell 167, 405–418 e413 (2016).

31. K. T. Roybal et al., Engineering T Cells with Customized Therapeutic Response Programs Using Synthetic Notch Receptors. Cell 167, 419–432 e416 (2016).

32. W. L. Chew et al., A multifunctional AAV-CRISPR-Cas9 and its host response. Nat Methods 13, 868–874 (2016).

33. A. M. Moreno et al., Immune-orthogonal orthologues of AAV capsids and of Cas9 circumvent the immune response to the administration of gene therapy. Nat Biomed Eng 3, 806–816 (2019).

34. C. H. Wang et al., Monitoring of the central blood pressure waveform via a conformal ultrasonic device. Nat Biomed Eng 2, 687–695 (2018).

35. E. S. Boyden, F. Zhang, E. Bamberg, G. Nagel, K. Deisseroth, Millisecond-timescale, genetically targeted optical control of neural activity. Nature neuroscience 8, 1263–1268 (2005).

36. B. Z. Fite et al., Magnetic resonance thermometry at 7T for real-time monitoring and correction of ultrasound induced mild hyperthermia. PloS one 7, e35509 (2012).

37. M. Otto-Duessel et al., In vivo testing of Renilla luciferase substrate analogs in an orthotopic murine model of human glioblastoma. Mol Imaging 5, 57–64 (2006).

38. M. W. McBurney et al., The mouse Pgk-1 gene promoter contains an upstream activator sequence. Nucleic Acids Res 19, 5755–5761 (1991).

