## Supplementary Materials for "Acoustogenetic Control of CAR T Cells via Focused Ultrasound"

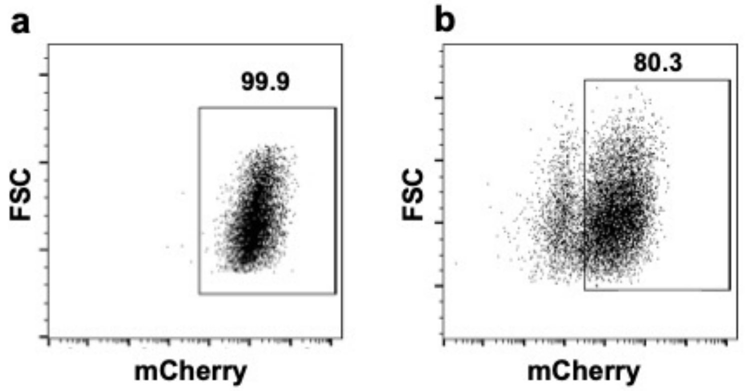


**Fig. S1. Transduction efficiencies of the dual-promoter eGFP reporter.** Representative flow cytometry data showing the lentiviral transduction efficiencies of the dual-promoter eGFP reporter (Hsp-eGFP-PGK-mCherry) in **(a)** HEK293T cells (99.9%) and **(b)** primary human T cells (80.3%). The mCherry+ gates are based on the profiles of the corresponding non-engineered cells.

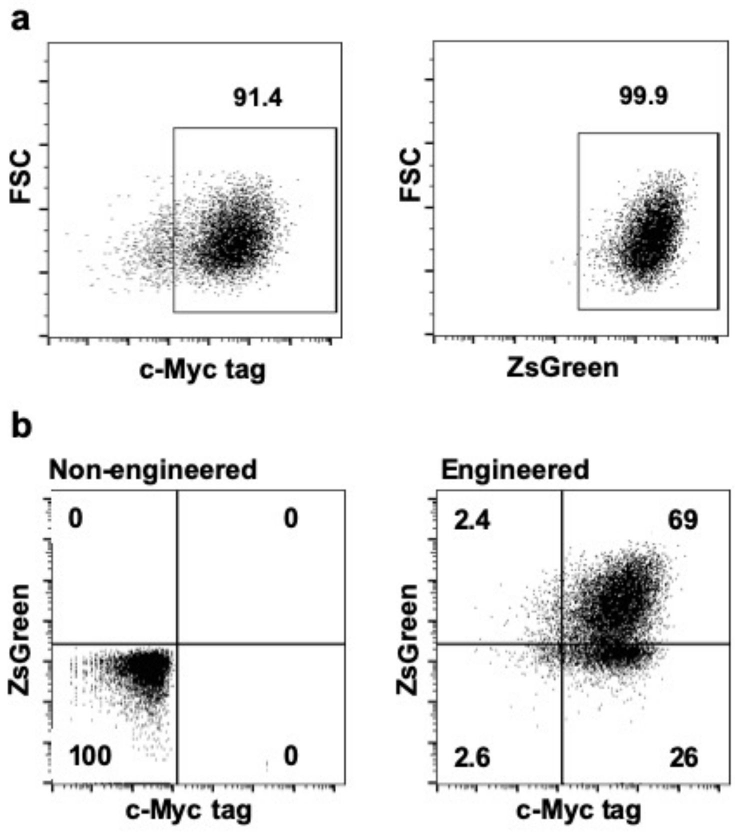


**Fig. S2. Transduction efficiencies of the heat-inducible Cre-lox system in Jurkat and primary human T cells. (a)** Jurkat cells were lentivirally co-transduced with inducible Cre (Hsp-Cre-PGK-cMyc-LaG17) and lox-stop CAR reporter (PGK-LoxH-ZsGreen-stop-LoxP-CAR). Shown are the transduction efficiencies of cells without HS. **(b)** Primary human T cells were co-transduced with inducible Cre (Hsp-Cre-PGK-cMyc-LaG17) and lox-stop CAR reporter (PGK-LoxH-ZsGreen-stop-LoxP-CAR) lentiviruses. The cells were sorted through MACS for c-Myc+ cells. The representative transduction efficiency after MACS is 69% for the c-Myc and ZsGreen double positive, 95% for the c-Myc+ and 71.4% for the ZsGreen+ cells. Gating is based on the corresponding non-engineered cells with the same c-Myc antibody staining.

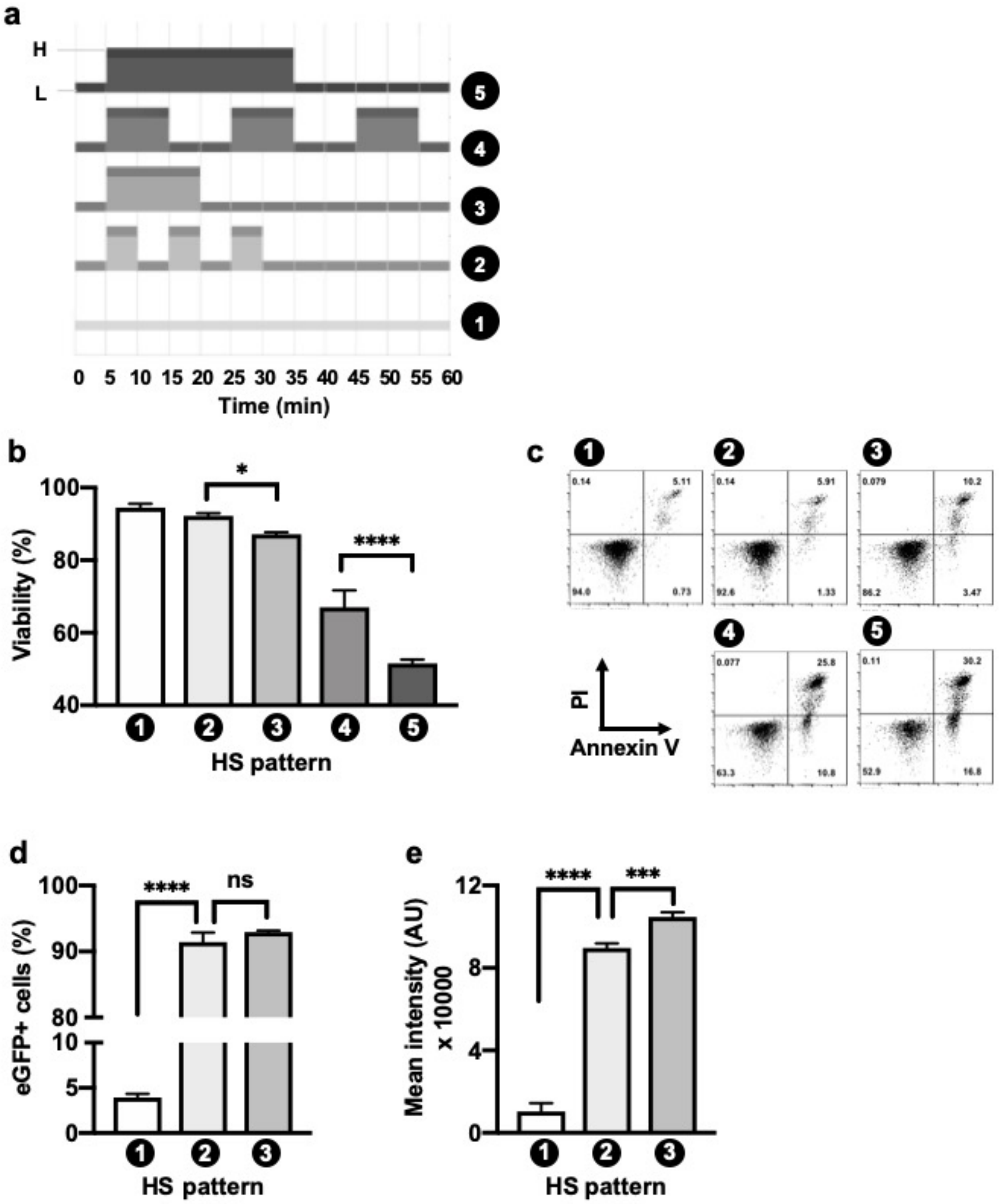
a
H
L
5
4
3
2
1
0 5 10 15 20 25 30 35 40 45 50 55 60
Time (min)
b
Viability (%)
100
80
60
40
*
****
1 2 3 4 5
HS pattern
c
1
0.14
5.11
94.0
0.73
2
0.14
5.91
92.6
1.33
3
0.079
10.2
86.2
3.47
4
0.077
25.8
63.3
10.8
5
0.11
30.2
52.9
16.8
PI
Annexin V
d
eGFP+ cells (%)
100
90
80
10
5
0
****
ns
1 2 3
HS pattern
e
Mean intensity (AU) x 10000
12
8
4
0
****
***
1 2 3
HS pattern

**Fig. S3. Thermal tolerance of primary human T cells. (a)** Different patterns of HS. H: heating at 43°C; L: resting at 37°C. **(b)** The viability of non-transduced primary human T cells 24 hr after different patterns of HS in (a) assayed by Annexin V and PI staining. **(c)** Representative flow cytometry data of (b). **(d-e) (d)** The percentage of eGFP+ cells and **(e)** their mean fluorescence intensity in primary human T cells engineered with the dual-promoter eGFP reporter in Fig. 1b under different HS patterns in (a). $N = 3$ repeats; error bar: SEM. *: $p < 0.05$; ***: $p < 0.001$; ****: $p < 0.0001$; ns: no significant difference.

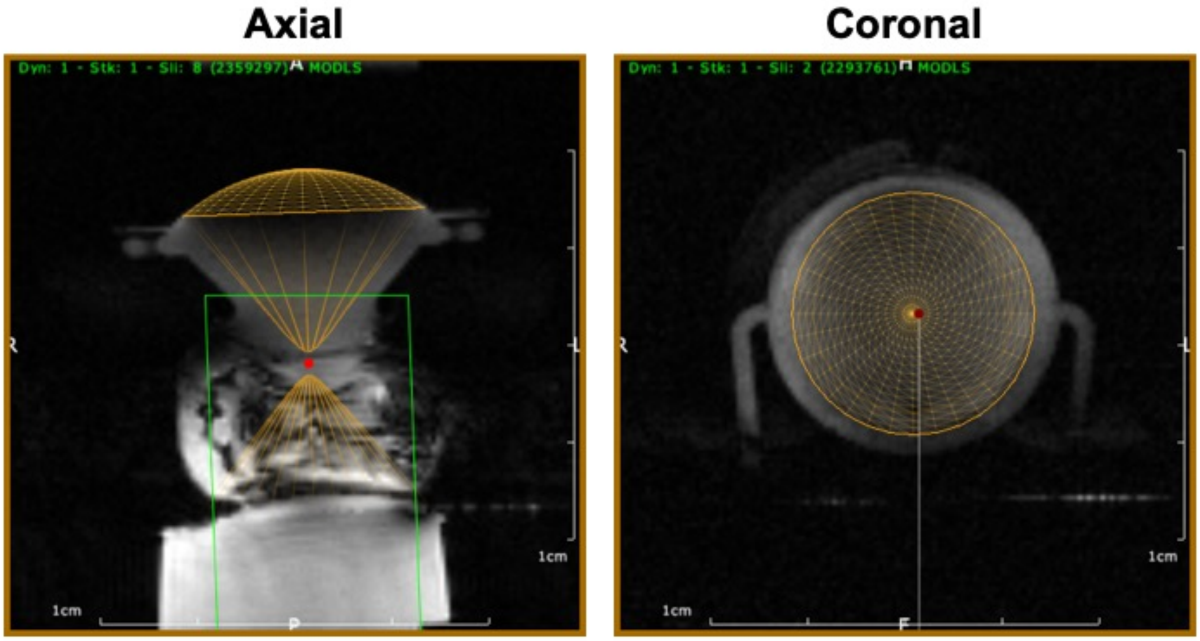


**Fig. S4. MRI images of the *in vivo* FUS experimental setup.** An anesthetized mouse is laid on its side on an MR bed containing an agarose gel pad. The ultrasound transducer is positioned above the targeted region on the mouse's hindlimb. Three-dimensional representations of the transducer generated by the ThermoGuide software are superimposed on the MRI images. Left: an axial view of the experimental setup. The bright red dot indicates the theoretical ultrasound focal point. Right: a coronal view showing the transducer.

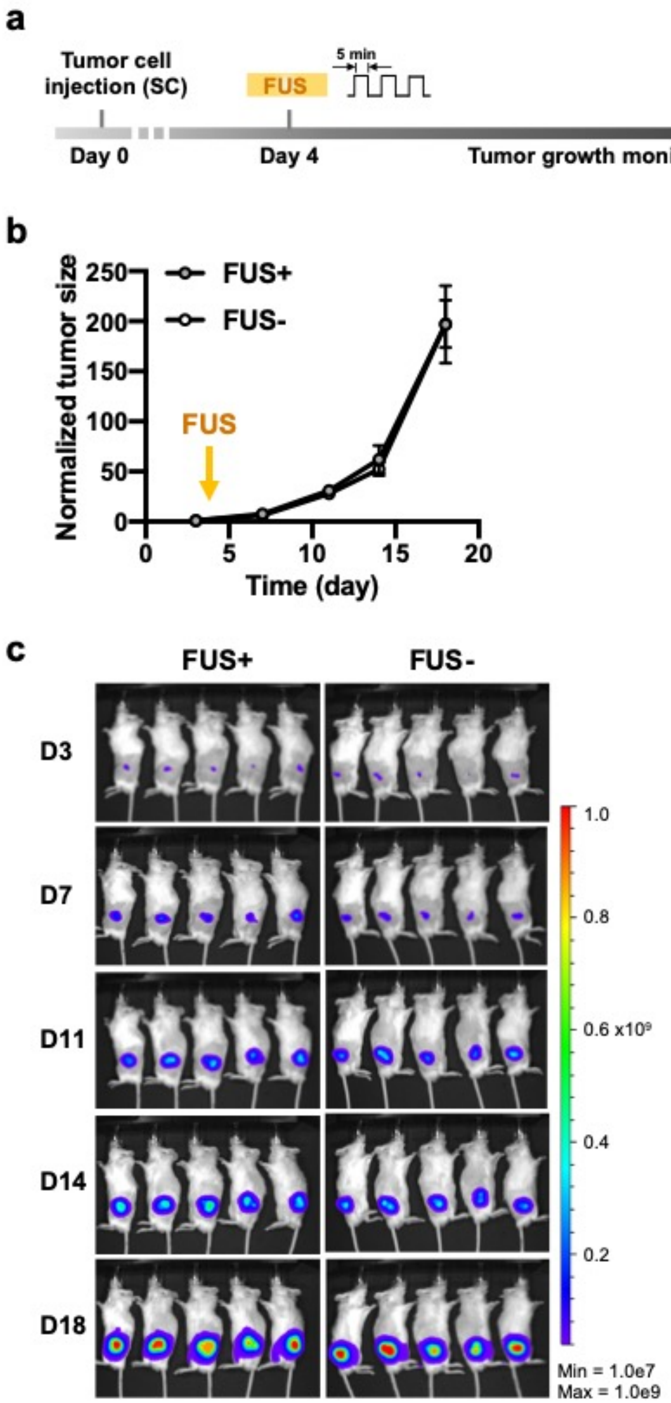

a
Tumor cell injection (SC)
FUS
5 min
Day 0
Day 4
Tumor growth monitoring
b
Normalized tumor size
FUS+
FUS-
FUS
Time (day)
c
FUS+
FUS-
D3
D7
D11
D14
D18
0.6 x10⁹
Min = 1.0e7
Max = 1.0e9

**Fig. S5. FUS has no detectable effect on tumor growth. (a)** Timeline of *in vivo* experiment. Fluc+ Nalm-6 cells are subcutaneously injected into NSG mice to generate matched bilateral tumors. Four days later, three pulses of 5-min FUS stimulation targeted at 43°C was applied on the tumor on one side (left), while the tumor on the other side (right) received no FUS stimulation. Tumor growth after FUS stimulation was monitored by bioluminescence imaging. **(b-c)** The **(b)** quantified tumor growth and **(c)** representative bioluminescence images of the tumors with (FUS+) or without (FUS-) FUS stimulation. Tumor size is quantified using the integrated luminescence intensity and normalized to that of the same tumor on the first measurement. N = 5 mice. Error bar: SEM. No significant difference was detected.

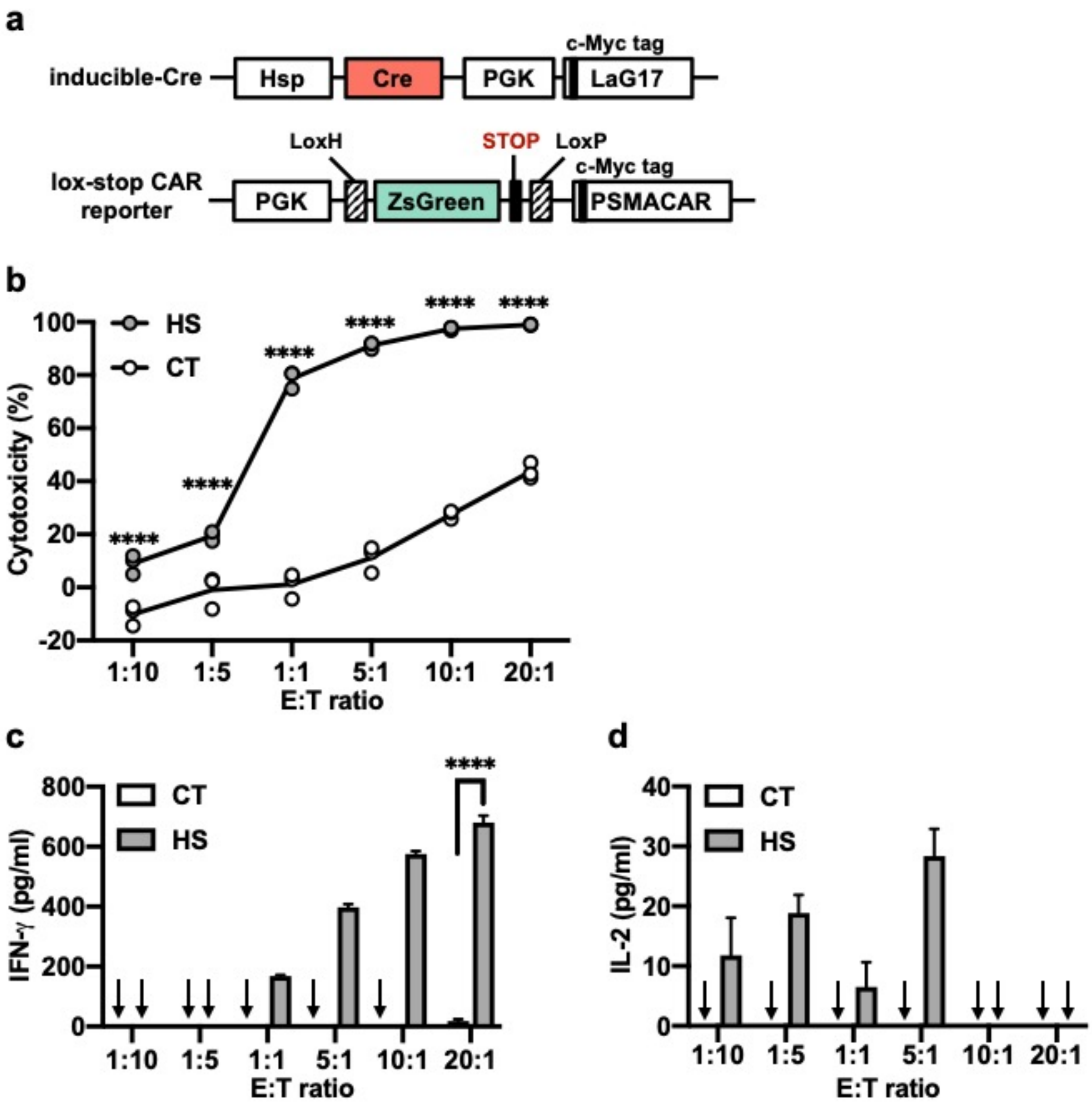


**Fig. S6. Functionality of the heat-inducible PSMACAR T cells *in vitro*. (a)** Schematics of the transgenes used in this figure, including the heat-inducible Cre and the lox-stop PSMACAR reporter. **(b)** Cytotoxicity of the T cells engineered with the transgenes in (a) against Fluc+ PSMA+ PC3 tumor cells at various E:T ratios. **(c-d)** Quantification of **(c)** IFN-γ and **(d)** IL-2 cytokine release associated with (b). Arrow: cytokine level not detectable. CT: without HS. HS: with a continuous 15-min HS. N = 3; error bar: SEM. ****: $p < 0.0001$.

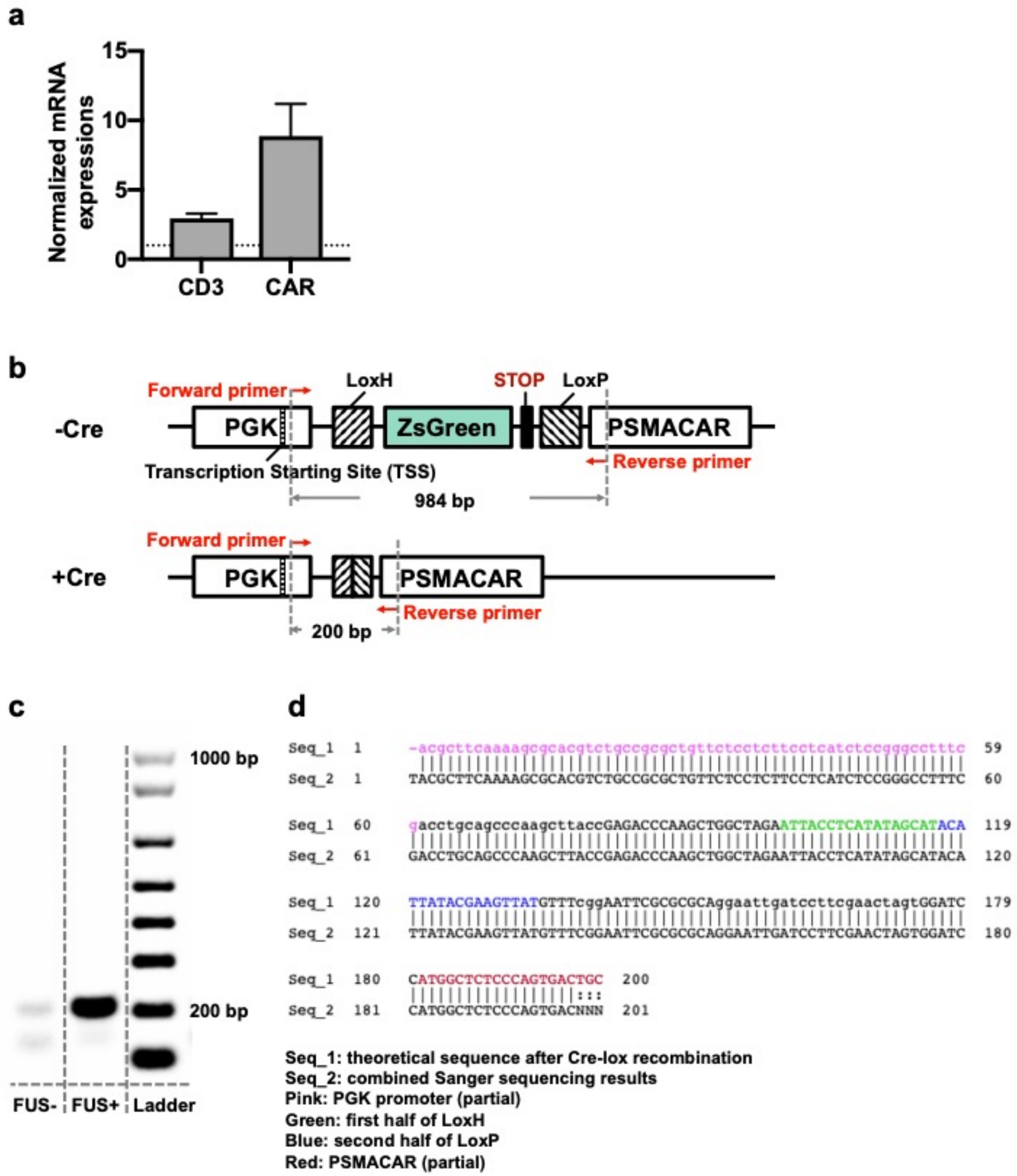
a
Normalized mRNA expressions
15
10
5
0
CD3
CAR
b
-Cre
Forward primer
LoxH
STOP
LoxP
PGK
ZsGreen
PSMACAR
Transcription Starting Site (TSS)
Reverse primer
984 bp
+Cre
Forward primer
PGK
PSMACAR
Reverse primer
200 bp
c
1000 bp
200 bp
FUS-
FUS+
Ladder
d
Seq_1 1 -acgcttcaaaagcgcacgtctgccgcgctgttctcctcttcctcatctccgggccttte 59
Seq_2 1 TACGCTTCAAAAGCGCACGTCTGCCGCGCTGTTCTCCTCTTCCTCATCTCCGGGCCTTTC 60
Seq_1 60 gacctgcagcccaagcttaccGAGACCCAAGCTGGCTAGAATTACCTCATATAGCATACA 119
Seq_2 61 GACCTGCAGCCCAAGCTTACCGAGACCCAAGCTGGCTAGAATTACCTCATATAGCATACA 120
Seq_1 120 TTATACGAAGTTATGTTTcggAATTCGCGCGCAggaattgatccttcgaactagtGGATC 179
Seq_2 121 TTATACGAAGTTATGTTTCGGAATTCGCGCGCAGGAATTGATCCTTCGAACTAGTGGATC 180
Seq_1 180 CATGGCTCTCCCAGTGACTGC 200
Seq_2 181 CATGGCTCTCCCAGTGACNNN 201
Seq_1: theoretical sequence after Cre-lox recombination
Seq_2: combined Sanger sequencing results
Pink: PGK promoter (partial)
Green: first half of LoxH
Blue: second half of LoxP
Red: PSMACAR (partial)

**Fig. S7. Quantification of relative mRNA expression levels in tumors. (a)** The expression levels of CD3 and CAR mRNA in tumors with FUS stimulation (Fig. 4E, FUS+), with their values normalized to the corresponding mRNA levels in tumors without FUS stimulation (Fig. 4e, FUS-). Dotted line: the corresponding mRNA levels in tumors without FUS stimulation are set to 1. **(b)** Schematics of the theoretical templates without (-Cre) or with (+Cre) Cre-lox recombination, and the expected products amplified by the designed primers. **(c)** Gel electrophoresis of the qPCR products of the tumors with FUS stimulation (FUS+) or without (FUS-). **(d)** Alignment of the combined Sanger sequencing results and the theoretical template with Cre-lox recombination.

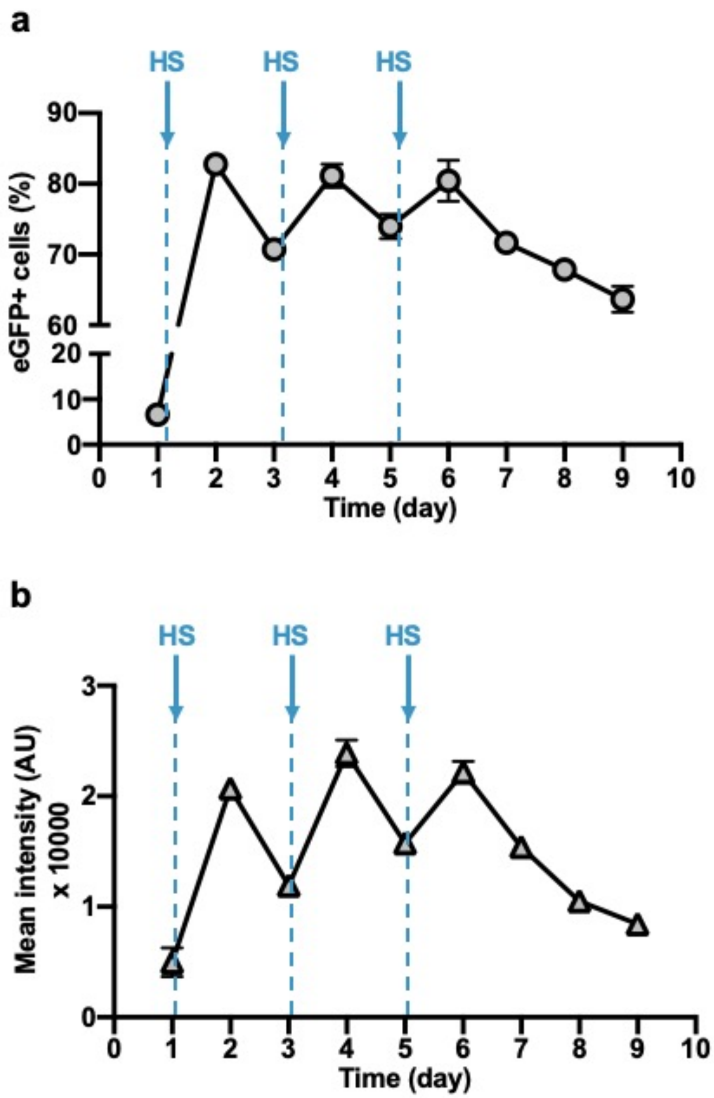


**Fig. S8. Gene activation with periodical HS. (a-b)** The dynamics of eGFP expression in terms of **(a)** percentage of eGFP+ cells and **(b)** mean eGFP intensity in primary human T cells transduced with the dual-promoter eGFP reporter and subjected to a 10-min HS every 48 hr on Day 1, 3, and 5. N = 3; error bar: SEM.

| Plasmid | Purpose | Used in | Source |
|---|---|---|---|
| pHR-Hsp-eGFP-PGK-mCherry | inducible GFP reporter | Figs. 1C-G, S1, S7 | This work |
| pHR-PGK-LoxH-ZsGreen-STOP-LoxP-cMyc-CD19CAR | Lox recombination CD19CAR reporter | Figs. 2B, 2D-I, 4B-C, S2 | This work |
| pHR-Hsp-Cre-PGK-cMyc-LaG17 | inducible Cre | Figs. 2B, 2D-I, 4B-C, S2, S5, S6 | This work |
| pHR-Hsp-Fluc-PGK-Rluc-P2A-mCherry | inducible dual luciferase reporter | Fig. 3D, 3G-H | This work |
| pHR-PGK-LoxH-ZsGreen-STOP-LoxP-cMyc-PSMACAR | Lox recombination PSMACAR reporter | Figs. 4D-E, S5, S6 | This work |
| pHR-PGK-PSMA | PSMA antigen | Figs. 4D-E, S5 | This work |
| pHIV-EF1a-Fluc-IRES-ZsGreen | firefly luciferase | Figs. 2G-I, 4B-E, S4, S5 | Addgene #39196 |

**Table S1. Plasmids used in this work.** Detailed information on the plasmids used in this work, including the features of the constructs, their usage, and sources.

| Pattern | Step | Temperature | Time |
|---|---|---|---|
| 1 | | 37°C | ∞ |
| 2 | Initial equilibrium | 37°C | 5 min |
| | 3 Cycles | 43°C | 5 min |
| | | 37°C | 5 min |
| 3 | Initial equilibrium | 37°C | 5 min |
| | 1 Cycle | 43°C | 15 min |
| 4 | Initial equilibrium | 37°C | 5 min |
| | 3 Cycles | 43°C | 10 min |
| | | 37°C | 10 min |
| 5 | Initial equilibrium | 37°C | 5 min |
| | 1 Cycle | 43°C | 30 min |

**Table S2. Thermal cycler HS patterns used in this work.** The patterns of different HS treatments performed in this work using a thermal cycler.

| Cell type | Gene delivery method | Construct | Sorting | Efficiency | Application |
|---|---|---|---|---|---|
| HEK 293T | Lentivirus | Hsp-eGFP-PGK-mCherry | Not sorted | >99% | Gene activation in vitro by HS |
| Jurkat | Lentivirus | Hsp-eGFP-PGK-mCherry | Not sorted | >99% | Gene activation in vitro by HS |
| Jurkat | Lentivirus | Hsp-Cre-PGK-cMyc-LaG17 | Not sorted | 91.40% | Cre activation in vitro by HS, CD69 assay |
| | (co-transduction) | PGK-LoxH-ZsGreen-STOP-LoxP-CD19CAR | Not sorted | >99% | |
| Nalm-6 | Lentivirus | Hsp-Fluc-PGK-Rluc-mCherry | FACS | >99% | Gene activation in vivo by FUS |
| Nalm-6 | Gift from Michel Sadelain Lab | constitutive Fluc+ | FACS | >99% | In vitro and in vivo cytotoxicity studies |
| PC3 | Lentivirus | pHIV-EF1a-Fluc-IRES-ZsGreen (Addgene) | FACS | >99% | In vitro and in vivo cytotoxicity studies |
| | (co-transduction) | PGK-PSMA | FACS | >99% | |

**Table S3. Engineered cells used in this work.** Detailed information on the engineered cells (excluding primary human T cells) used in this work, including the cell type, the transgene delivered and the associated delivery method and efficiency, sorting information and applications.

**Movie S1. The dynamics of HS-activated eGFP expression.** Real-time fluorescence imaging of two HEK 293T cells transduced with the dual-promoter eGFP reporter (Hsp-eGFP-PGK-mCherry). A 15-min HS at 43°C was applied from 30 to 45 min after the initiation of imaging using a heating stage integrated with the microscope. Imaging lasted for 12 hours. GFP: GFP channel showing the dynamic activation of eGFP; mCherry: mCherry channel showing the constitutive mCherry expression.

**Movie S2. MRI-guided FUS stimulation on a targeted region on the hindlimb of an anesthetized mouse.** MRI images are superimposed with color-coded temperature map of the region of interest. FUS stimulation was aimed at the hindlimb of an anesthetized mouse. The FUS stimulation is composed of a temperature rising phase (Dyn 1 - 10), a steady-state phase (Dyn 11 - 75), and a cool-down phase (Dyn 76-90). Dyn: dynamic; one dynamic of MRI scanning takes 4.6 sec.
